# Development of Patient-Derived Neuroprogenitor Cells (hNPCs), Neurons and Astrocytes to Explore the Etiology of Guam Parkinsonism-Dementia Complex (PDC)

**DOI:** 10.1101/2025.10.07.680309

**Authors:** Anna C. Chlebowski, Y. Yang, Nailah A. Siddique, Teepu Siddique, Peter S. Spencer, John C. Steele, Glen E. Kisby

## Abstract

Parkinsonism-Dementia Complex (PDC) is one phenotype of a disappearing neurodegenerative disease (Guam ALS-PDC) that shows clinical and neuropathological relationships with amyotrophic lateral sclerosis (ALS), atypical parkinsonism and Alzheimer’s disease. ALS-PDC has been linked with exposure to environmental factors (notably cycad plant neurotoxins), but evidence from human and animal studies is inconclusive. Patient-derived induced pluripotent stem cells (iPSCs) provide a powerful in vitro system to explore the underlying cause of PDC. iPSC lines were derived from lymphocytes of a PDC-affected Guamanian Chamorro female patient and an age- and gender-matched healthy Chamorro resident of PDC-unaffected Saipan using non-integrating episomal plasmids. iPSCs derived from both patients expressed pluripotency markers (Oct4, SSEA-4, TRA-1-60, Sox2) prior to the generation of neuroprogenitor cells (hNPCs), neurons and astrocytes. An embryoid body protocol was used to derive hNPCs from both iPSC lines while a differentiation media was used to generate neurons from hNPCs. hNPCs derived from both iPSC patients’ lines displayed established neuroprogenitor markers (nestin, Sox2), while the differentiated hNPCs exhibited both neuronal (beta-tubulin III, Map2, doublecortin) and synaptic (synaptophysin, PSD-95) markers. Expression of these protein markers in hNPCs and neurons by dot blotting was also observed for both lines. Astrocyte progenitor cells and mature astrocytes with appropriate markers were also developed from the hNPCs of both lines using commercial kits. Development of these patient-derived iPSCs provides a human model for evaluating the role of environmental (e.g., cycad toxins) and genetic factors in ALS-PDC and possibly other related neurodegenerative diseases.

## Introduction

Western Pacific Amyotrophic Lateral Sclerosis and Parkinsonism-Dementia Complex (ALS-PDC) is a progressive neurodegenerative disease with multiple clinical phenotypes (ALS, atypical parkinsonism with dementia (P-D), dementia alone) and of familial or sporadic origin that was formerly highly prevalent among the island communities of the southern Marianas (Guam and Rota), Honshu, Japan (Kii Peninsula), and New Guinea (Papua, Indonesia). Following WWII, the prevalence of ALS, and later of P-D, declined in Guam, Kii-Japan and Papua, Indonesia, such that the disease is predicted to soon disappear from all three geographic isolates. The disappearance of an hyperendemic disease among these diverse ethnic genotypes suggests the operation of one or more common environmental factors to which the affected populations were formerly exposed (Gimenez-Roldan et al. 2021; Spencer et al. 2016). Leading etiological candidates for ALS/PDC are the cycad plant toxins [cycasin, methylazoxymethanol (MAM) and β-*N-*methylamino-L-alanine (L-BMAA)] present in traditional food and/or medicine (Kisby and Spencer 2011; Spencer et al. 2016; 2020). While the cause of this prototypical neurodegenerative disorder is most probably attributable to cycad toxins, evidence from human and animal studies remains inconclusive.

Induced pluripotent stem cells (iPSCs) are an important tool for uncovering disease mechanisms and developing therapeutic strategies to treat them (Li et al. 2018; Pazzin et al. 2024; Rivetti di Val Cervo et al. 2021). iPSCs have also emerged as a powerful model for the study of neurodegenerative diseases, including ALS, Parkinson disease and Alzheimer disease (AD) (Amponsah et al. 2021; Ceccarelli et al. 2024; Okano and Morimoto 2022; Riemens et al. 2020; Samson et al. 2024; Valadez-Barba et al. 2021). iPSC-derived neural cell models from AD patients have allowed for modeling of tau and β-amyloid pathologies beyond what was possible with animal models (Choi et al. 2015; Choi et al. 2014; Choi et al. 2016). iPSC-derived neurons from both familial and sporadic AD-diseased patients express significantly higher levels of pathological proteins (e.g., amyloid, phosphotau isoforms) relative to controls (Israel et al. 2012). iPSC-derived astrocytes have also revealed an important role in the pathogenesis of AD (Lee et al. 2025) and ALS/PDC in Kii-Japan (Kobayashi et al. 2023; Leventoux et al. 2024). Moreover, long-term culture of iPSCs from AD patients produces organoids with pathological features similar to those observed in the AD brain (Karmirian et al. 2023; Kim et al. 2024; Marei et al. 2023). Thus, iPSCs may also be helpful in elucidating the environmental and any genetic factors that contribute to the pathogenesis of ALS/PDC, a prototypical neurodegenerative disorder characterized by a tau-dominated polyproteinopathy (Forman et al. 2002; Gimenez-Roldan et al. 2021; Miklossy et al. 2008; Mimuro et al. 2018; Sebeo et al. 2004; Smith et al. 2024; Verheijen et al. 2020; Winton et al. 2006).

The primary somatic cell type of choice for generating iPSCs was formerly dermal fibroblasts (Takahashi et al. 2007; Yu et al. 2007), but other tissue sources, including lymphoblastoid cell lines (LCLs), have recently become a rich source because they usually contain extensive donor information with regard to genotype and existing genomic and phenotypic data (Gurwitz and Steeg 2024; Kumar et al. 2020). While primary cells from the skin (i.e., fibroblasts) and blood (e.g., lymphocytes) of patients with neurodegenerative disease are routinely used, improvements in the reprogramming efficiency of immortalized lymphoblastoid cells from donors with neurodegenerative disease provides another source of cells for understanding the cellular and molecular mechanisms of these brain disorders (Barrett et al. 2014; Hedges et al. 2021; Kumar et al. 2020; Kumar et al. 2016; Walker et al. 2021). Lymphoblastoid cells (LCLs) are generated by infecting peripheral blood lymphocytes with the Epstein Barr Virus (EBV) (Sie et al. 2009). The reprogramming of LCLs from donors with various neurological disorders into iPSCs has become highly efficient such that they are functionally indistinguishable from fibroblast-derived iPSCs (Barrett et al. 2014; Fujimori et al. 2016; Kumar et al. 2020) and formed without the reprogramming of viral elements (Chapotte-Baldacci et al. 2023; Rajesh et al. 2011). The present study utilizes LCLs generated from Guam Chamorro PDC-affected and PDC-free Saipan Chamorro control patients to derive iPSCs, neuroprogenitor cells (hNPCs), neurons and astrocytes to evaluate the neurotoxic properties of suspected environmental factors, notably cycad neurotoxins (Gimenez-Roldan et al. 2021; Spencer 2022). The LCL lines were previously developed by the late Dr. John Steele and Dr. Teepu Siddique to evaluate the occurrence of genetic mutations in Guam familial PDC cases, but failed to identify a single mutant gene locus (Morris et al. 2004). Familial ALS-PDC likely arises from transgenerational exposure to cycad toxins (Spencer 2020). Development of iPSCs from LCLs could provide important information about the complex etiology of ALS/PDC and possibly other related neurodegenerative diseases (Spencer et al. 2016; Steele et al. 2015).

## Materials and Methods

### 2.1 Lymphocytes

Lymphocytes from Chamorros living on Guam (diagnosed with PDC) or Saipan (normal control) with detailed patient histories and genetic information were immortalized with EBV to lymphoblastoid cell lines as previously described (Morris et al. 2004; Steele et al. 2015). Of the 90 lymphoblastoid cell lines (LCLs) that were successfully developed (Morris et al. 2004), 10 pairs of age- and sex-matched LCLs were selected from both the Guamanian and the Saipanese Chamorro patients. Saipan was chosen to develop healthy control LCL because subjects had no history of cycad consumption since the trees had been removed from the island in the 1920s in order to grow sugarcane (McGeer and Steele 2011; Zhang et al. 1990). The clinically normal Chamorro from Saipan (female, age 40) was born on-island in 1959 while the Guam-born PDC patient (female, age 57) had advanced brain disease (J. Steele, *personnel communication*). LCLs from all pairs of lines were expanded by culturing the cells in RPMI media (Gibco, Thermofisher Scientific, Waltham, MA) that contained 15% FBS (Hyclone, Marlborough, MA) and 100 units/mL pen/step (Gibco). Cultures were maintained at 37°C with 5% CO_2_ and monitored daily for media changes and the formation of aggregates. LCLs from one pair of age- and sex-matched Saipan control and Guam PDC subjects were selected for reprogramming to induced pluripotent cells (iPSC) according to the protocols of Barrett and colleagues (Barrett et al. 2014).

### 2.2 LCL Transfection

LCLs from the healthy control and PDC patient were transfected using episomal plasmids expressing pluripotency factors and p53 shRNA in combination with small molecules according to the user manual for the Neon^®^ Transfection Instrument (ThermoFisher, Waltham MA). Non-integrating episomal plasmids (Trevisan et al. 2017) for hSK, hUL, hOCT3.4, ET2K were obtained from AddGene (Watertown, MA), and the plasmids were prepared and purified by GeneWiz (Plainfield, NJ) for LCL transfection (**Supplement 1**). Three separate transfection reactions (pulsed 2 x 30 ms at 1100 volts) were carried out for each line on two separate days, using 1×10^5^ cells and 1.0 μg of each plasmid in a single reaction. The transfected LCLs were then transferred to 24-well Matrigel^®^ (Corning, NY)-coated plates before transitioning them from LCL culture media to reprogramming media over the next several days. After 2-3 weeks, the transfected LCLs were gradually transitioned to mTeSR^®^-1 media (STEMCELL^®^ Technologies, Vancouver, BC), with iPSC colonies appearing over the next several weeks. iPSCs from the reference line NCRM5 (ND50031) were obtained from RUCDR Infinite Biologics (Piscataway, NJ) and cultured in a similar fashion as the Chamorro iPSC lines. This reference line differentiates into hNPCs, neurons, and astrocytes (Malik et al. 2014; Pandya et al. 2017) and was used to develop the optimal protocol for the development of hNPCs and neurons from the PDC-affected and control Chamorro iPSC lines.

### 2.3 Induced pluripotent stem cells (iPSCs)

iPSC colonies were manually subcloned into Matrigel^®^ pre-coated plates. Areas of differentiation were manually removed and desirable colonies manually sub-cloned to obtain a pure population of iPSCs. When the iPSC colonies were < 10% differentiated, they were treated with EZ-LiFT™ reagent (Millipore Sigma, Burlington, MA) and passed for a minimum of 10 passages before neural induction. Both iPSC lines were characterized for pluripotency by immunocytochemistry at passage 3-5 and again at passage 10 before neural induction. iPSCs were cultured at 37°C with 5% CO_2_ with daily medium changes. Genomic stability of the two iPSC lines and the corresponding LCLs was assessed using the KaryoStat™ Assay (ThermoFisher) (MacArthur et al. 2019; Ramme et al. 2021).

### 2.4 Neural Progenitors (hNPCs)

The STEMdiff™ Embryoid Body (EB) protocol with SMADi (STEMCELL™ Technologies) was used to generate neural progenitor cells (hNPCs) from iPSCs. This method was optimal for producing hNPCs from the two Chamorro iPSC lines (**Supplement 2**). hNPCs were maintained by daily media changes at 37°C with 5% CO_2_ and the cells passed every 4-6 days. At passage 2-3, hNPCs were fixed with 4% buffered paraformaldehyde (PFA) for characterization of markers by immunocytochemistry.

### 2.5 Neural Differentiation

The BrainPhys™ neuronal medium hPSC neuron kit (STEMCELL™ Technologies, Seattle WA) was determined to be optimal for generating neurons from Chamorro-derived hNPCs (**Supplement 3**). hNPCs (passage 3-4) were seeded at a density of 40,000 - 60,000 cells/cm^2^ in poly-L-ornithine/laminin-coated 96-well plates and 24h later transitioned to BrainPhys™ Neuronal Medium (STEMCELL™ Technologies) containing NeuroCult™ SM1 Supplement (STEMCELL™ Technologies), 1x N2 Supplement A (STEMCELL™ Technologies), 20 ng/mL BDNF (Peprotech, Cranbury, NJ), 20 ng/mL GDNF (Peprotech), 1 mM dibutyryl c-AMP (STEMCELL™ Technologies), and 200 nM ascorbic acid (Sigma, St Louis, MO). Cells were maintained with half-media changes every 2-3 days. On day 21 or 28 of differentiation, the cells were fixed with PFA for characterization of neuronal markers by immunocytochemistry.

#### Astrocyte progenitors and mature astrocytes

Mature astrocytes (ax0665) were obtained from Axol Bioscience (Cambridge, UK) and cultured according to the manufacturer’s protocol. The STEMdiff™ Astrocyte Differentiation and Maturation kits (STEMCELL™ Technologies) were used for the generation of astrocyte progenitors and mature astrocytes from both Chamorro-derived iPSC lines. The first 12 days of the protocol followed the STEMdiff™ Embryoid Body (EB) Protocol +SMADi (STEMCELL™ Technologies) for the generation of hNPCs from iPSCs (as described above). Following rosette selection, hNPCs were transitioned to astrocyte differentiation medium for 3 weeks to generate astrocyte progenitor cells (APCs). After the third week, astrocyte progenitors were characterization by immunocytochemistry or further differentiated for 3 weeks using astrocyte maturation media to generate mature astrocytes. Astrocytes, which were are considered mature as per the Kit instructions, were collected for final characterization.

### 2.6 Immunocytochemistry

iPSCs were fixed with 4% buffered PFA for 20-30 min before they were examined for pluripotency markers (SSEA-4, OCT-4, SOX2, and TRA-1-60) using the PSC 4-Marker Characterization Kit (Invitrogen, Waltham, MA). iPSC-derived NPCs, neurons and astrocytes were similarly fixed with 4% buffered PFA to immunoprobe for markers. Details on the method and markers used are given in **Supplement 4**. Immunoprobed iPSCs, hNPCs, neurons and astrocytes were imaged with a Keyence BZ-X710 Epifluorescence microscope equipped with the appropriate filters.

### 2.7 Dot blotting

Protein extracts of iPSC-derived hNPCs and neurons were examined for several markers by dot blotting (Chlebowski and Kisby 2020). Briefly, hNPCs (∼1.9 x 10^6^) from each iPSC line were seeded into 6-well plates coated with poly-L-ornithine and laminin; cells were collected after 24 h in media (hNPCs) or after differentiation with BrainPhys™ media for 28 days (neurons), as noted above. hNPCs and neurons from both lines were re-suspended in buffer containing cOmplete™ Mini EDTA-free protease inhibitor (Roche, Basel, CH), the cells sonicated on ice (Qsonica, Newton, CT), the homogenates centrifuged, and the supernatant analyzed for protein content (Bradford assay, BioRad, City) before storing at −80°C. Detailed information on the dot blotting of hNPCs and neurons is given in **Supplement 6**.

### 2.8 Statistics

Relative intensity values of each sample are presented as mean (n=2) ± standard deviation. Groups were compared using a two-way ANOVA with significance set at *p*<0.05. GraphPad Prism 8 was used for statistical analysis and the generation of graphs.

## Results

### 3.1 Generation and Characterization of iPSCs

Subsets of LCL lines (n=10) from healthy controls and PDC Chamorros (Steele et al. 2015) were selected for the development of iPSCs. LCLs from the healthy Saipan control and Guam PDC patient were first used to generate iPSCs using a previously published protocol (Barrett et al. 2014). iPSCs from both LCL lines exhibited similar morphology, as well as the expression of pluripotency markers (**Figure 1A**). The iPSC line derived from the Saipan subject grew in discrete colonies with well-defined edges, whereas the Guam PDC colonies showed less-defined edges, and less uniform expression of pluripotency markers in individual colonies, which is consistent with that reported for iPSCs developed from PDC patients from Kii Penninsula of Japan (Kobayashi et al. 2023). Both lines expressed all four pluripotency markers (SSEA4, OCT4, Sox2, TRA-1-60), which is consistent with that reported for the development of iPSCs from LCLs of patients with spinal muscular atrophy and sporadic and familial Parkinson disease (Barrett et al. 2014; Fujimori et al. 2016; Kumar et al. 2016). Karyotyping of both iPSC lines showed that they displayed a normal karyotype (**Figure 1B**). Karyotyping of the parental LCL lines also revealed a normal genotype (**Supplement 1**). Karyotyping of LCLs from two other pairs of healthy and PDC patients also showed a normal karyotype (*data not shown*).

**Figure 1.**
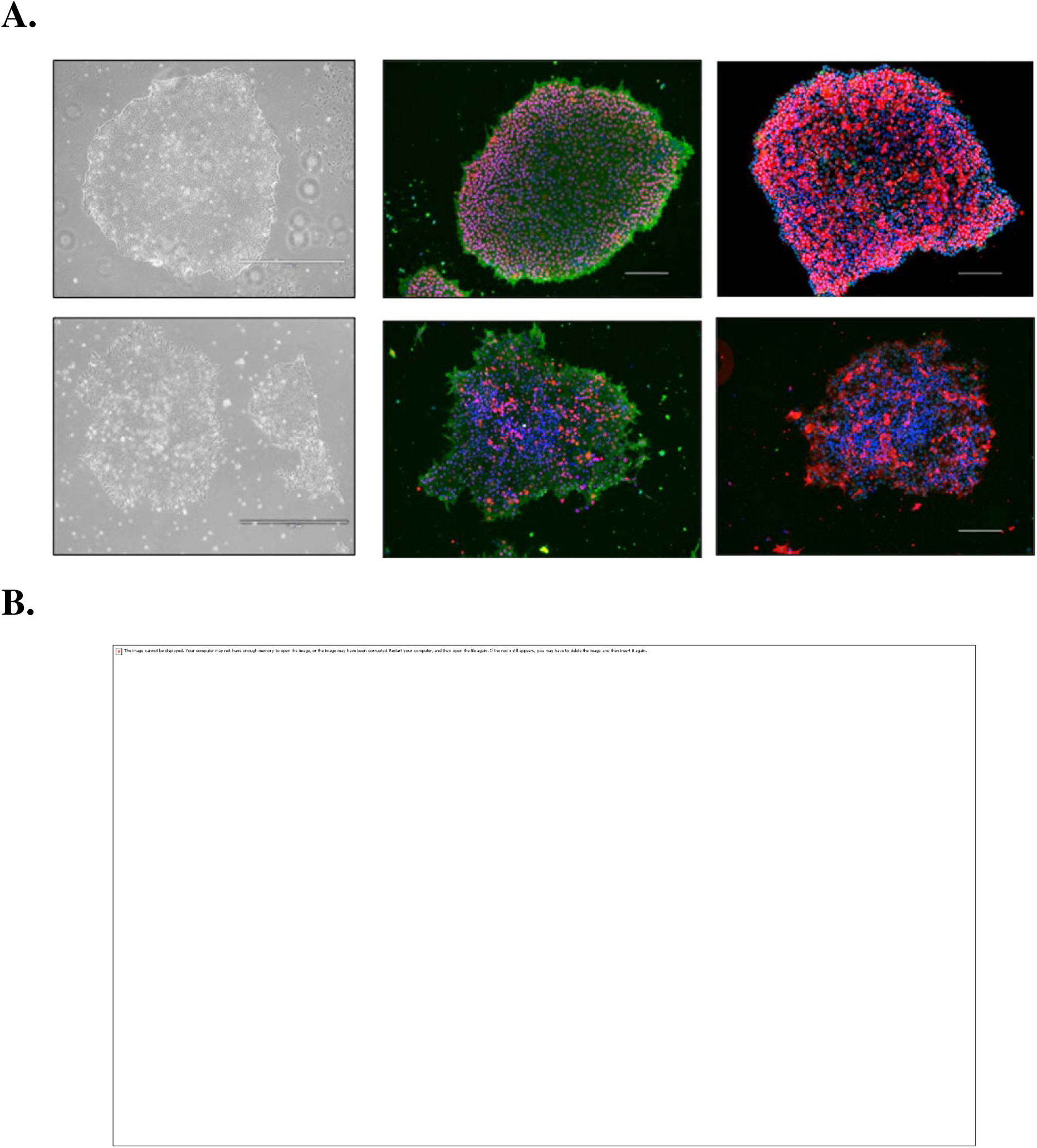
Characterization of Chamorro-derived iPSC lines. **A.** iPSCs were cultured using the matrigel/mTeSR-1 culture system for a minimum of 10 passages prior to their characterization and induction. iPSCs from the healthy control Saipan subject (*top row*) and Guam PDC patient (*bottom row*) were immunoprobed for markers of pluripotency (SSEA-4, OCT4, Sox2, TRA-1-60; *green or red*) and counterstained with DAPI (*blue*). **B.** Karyostat analysis of corresponding iPSC lines from the Saipan subject (*top*) and Guam (*bottom*) patient. The whole-genome view displays all somatic and sex chromosomes in one frame with high-level copy number. The smooth signal plot (right y-axis) is the smoothing of the log2 ratios that depict the signal intensities of probes on the microarray. A value of 2 represents a normal copy number state (CN = 2). A value of 3 represents chromosomal gain (CN = 3). A value of 1 represents a chromosomal loss (CN = 1). The pink, green and yellow colors indicate the raw signal for each individual chromosome probe, while the blue signal represents the normalized probe signal which is used to identify copy number and aberrations.

### 3.2 hNPC Generation and Characterization

iPSCs from both Chamorro subjects were induced to hNPCs using the STEMdiff^®^ Embryoid Body protocol. This method was selected from three different protocols that were used to induce the iPSC reference line NCRM5 into hNPCs (**Supplement 2**). hNPCs derived from both Chamorro iPSC lines were positive for the expected neuroprogenitor markers (nestin, Sox2) and negative for both pluripotent (SSEA4, OCT4) and differentiation markers (β-tubulin III, GFAP) (**Figure 2**). hNPCs from the Saipan healthy control also showed a morphology comparable to that of hNPCs derived from the reference line (**Supplement 2**).

**Figure 2.**
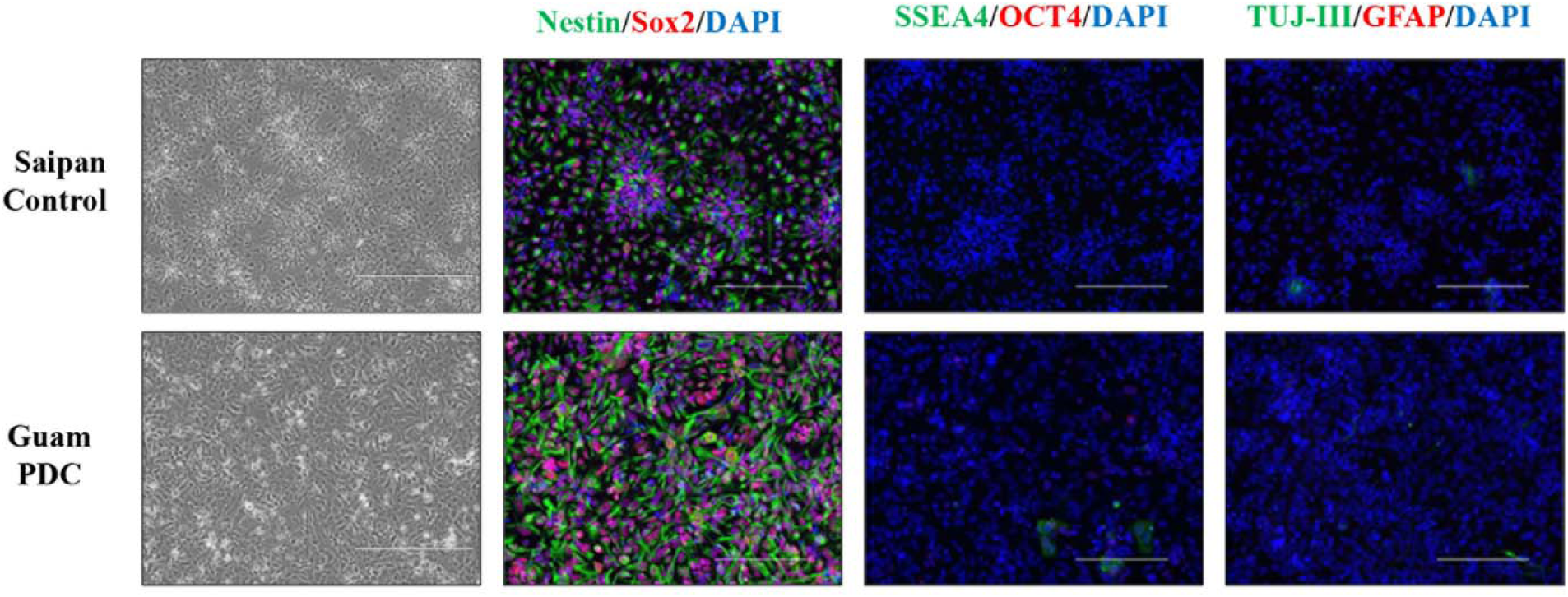
Characterization of Chamorro-derived hNPCs. hNPCs generated using the STEMdiff embryoid body protocol were collected and immunostained following completion of the protocol and grown for two subsequent passages in hNPC medium. hNPCs from both patient-derived lines were positive for hNPC markers (nestin, Sox2), and negative for markers of pluripotency (SSEA-4, OCT4) and differentiation (β-III-tubulin and GFAP). Cells were counterstained with DAPI nuclear stain.

### 3.3 Neuronal Differentiation

The optimal method for differentiating the iPSC-derived hNPCs into neurons was also determined from the differention of the NCRM5-derived hNPCs for 2-3 weeks using three different protocols (**Supplement 3**). The STEMdiff^®^ BrainPhys media protocol was used to differentiate the Chamorro-derived hNPCs because this method produced neurons with low staining for hNPCs and abundant staining for neuronal markers (β-tubulin III, doublecortin, MAP2) (**Supplement 3**). Differentiation of both Chamorro-derived hNPCs using this protocol produced cells expressing neuronal markers (β-tubulin III, doublecortin, MAP2) with little to no expresson of neuroprogenitor (nestin/SOX2) or glial markers (GFAP) (**Figure 3**). Moreover, the low expression of dendritic (MAP 2) and synaptic markers (synaptophysin, PSD-95) indicated that the differentiation of hNPCs for 21-28 days produced predominantly immature neurons. Expression of dendritic (MAP2) and synaptic markers (synaptophysin, PSD-95) also appeared to be lower in PDC-derived neurons than those from the control Chamorro (**Figure 3B**). These findings suggest that PDC-derived neurons were more immature than those derived from the healthy control patient. In support, PDC neurons also tended to migrate more than neurons derived from the Saipan control.

**Figure 3.**
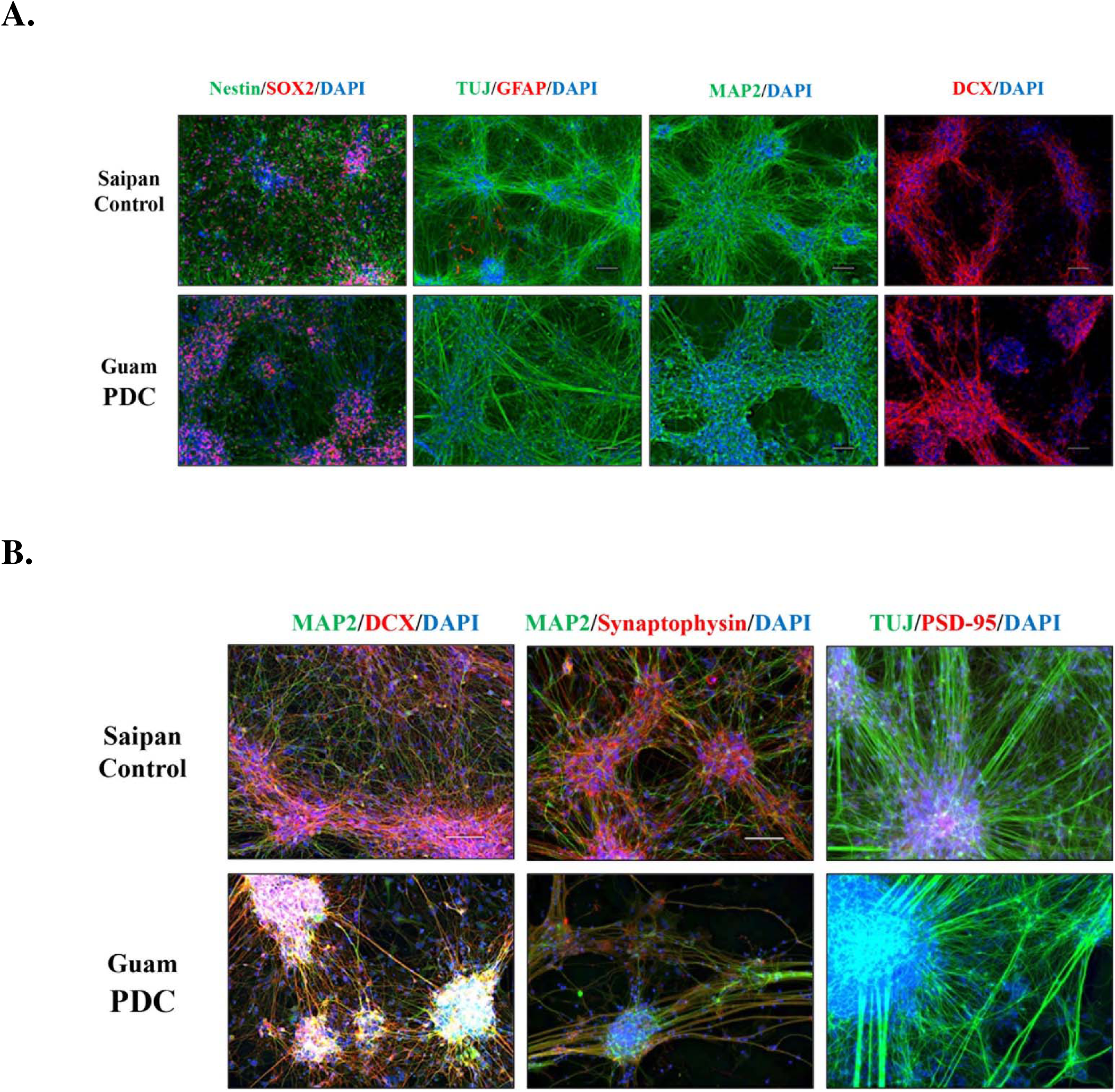
Characterization of Chamorro-derived neurons. Patient-derived hNPCs were differentiated into neurons using the STEMcell BrainPhys Media Kit protocol for 21 days (**A**) or 28 days (**B**) and the cells examined for markers of neural progenitors (nestin, Sox2), neurons (β-III-tubulin, Map2, DCX), glia (GFAP), and synapses (synaptophysin, PSD-95) and counterstained with DAPI.

The differentiated hNPCs from both Chamorro subjects were evaluated for the expression of neuronal and glial markers by dot-blot analysis, a rapid technique for determining differences in the expression of several neural markers (Chlebowski and Kisby 2020; Wehr and Levine 2012) (**Figure 4**). Neuronal markers were consistently higher in differentiated hNPCs than in corresponding hNPCs, an indication that the hNPCs were differentiated into neurons. Several neuronal markers were also higher in healthy Saipan Chamorro neurons than those derived from the Guam PDC subject. However, expression of nestin (neuroepithelial stem cell protein) was significantly higher (*p<0.05*) in PDC than in control neurons. β-tubulin III and doublecortin (DCX) were significantly higher (*p<0.05*) in the control than in the PDC line, and MAP2 also tended to be higher. Moreover, the levels of these markers in PDC neurons were ∼ 75% of that observed in control neurons. The glia marker GFAP was also significantly higher (*p<0.05*) in control than PDC neurons, suggesting that the glial cell population was also lower in PDC neuronal cultures. Expression of the presynaptic marker synaptophysin in the control line was ∼2x higher than the PDC line (p<0.05), while the postsynaptic marker PSD-95 also tended to be higher in control than PDC neurons. The differences in the expression of neuronal, glial and synaptic proteins suggested that control neurons were relatively more mature than PDC neurons. The higher expression of nestin in neurons derived from the PDC patient is also consistent with the delayed differentiation of hNPCs into neurons.

**Figure 4.**
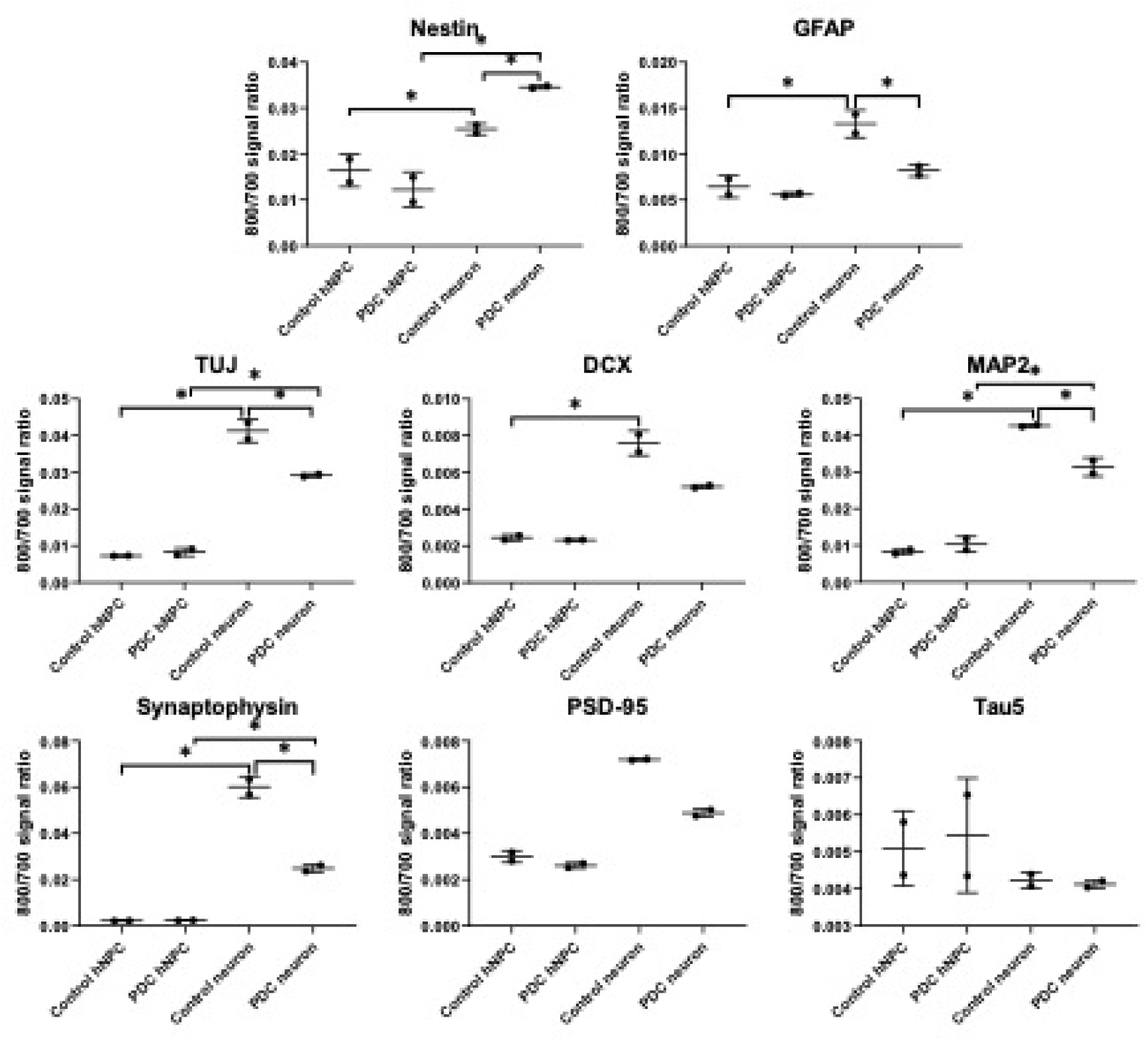
Level of protein markers in hNPCs and neurons from Chamorro subjects. hNPCs and neurons that had been differentiated for 28 days from a healthy Saipan control and Guam PDC patient were lysed and the homogenates analyzed by dot blot. Protein (1 µg) from hNPCs and neurons was applied to nitrocellulose membranes and then probed with antibodies to makrers for hNPCs (nestin), glia (GFAP), neurons (TUJ, DCX, MAP2,Tau 5), and synapse (synaptophysin, PSD-95). Values represent the ratio of the fluorescence intensity of the marker to that of total protein. *Significantly different (*p*<0.05) between hNPCs and neurons for the two cell lines by two way ANOVA.

### 3.4 Astrocyte progenitors and mature astrocytes

Astrocyte progenitors from both hNPC lines were positive for nestin and GFAP and negative for pluripotency markers (SSEA4, Oct4, Sox2, TRA-1-60) and β-tubulin expression indicating differentiaton to an astrocytic lineage (**Figure 5**). The astrocyte marker GFAP was also more prominent in control than PDC astrocyte progenitors suggesting that the glial cell population was lower in PDC cultures.

**Figure 5.**
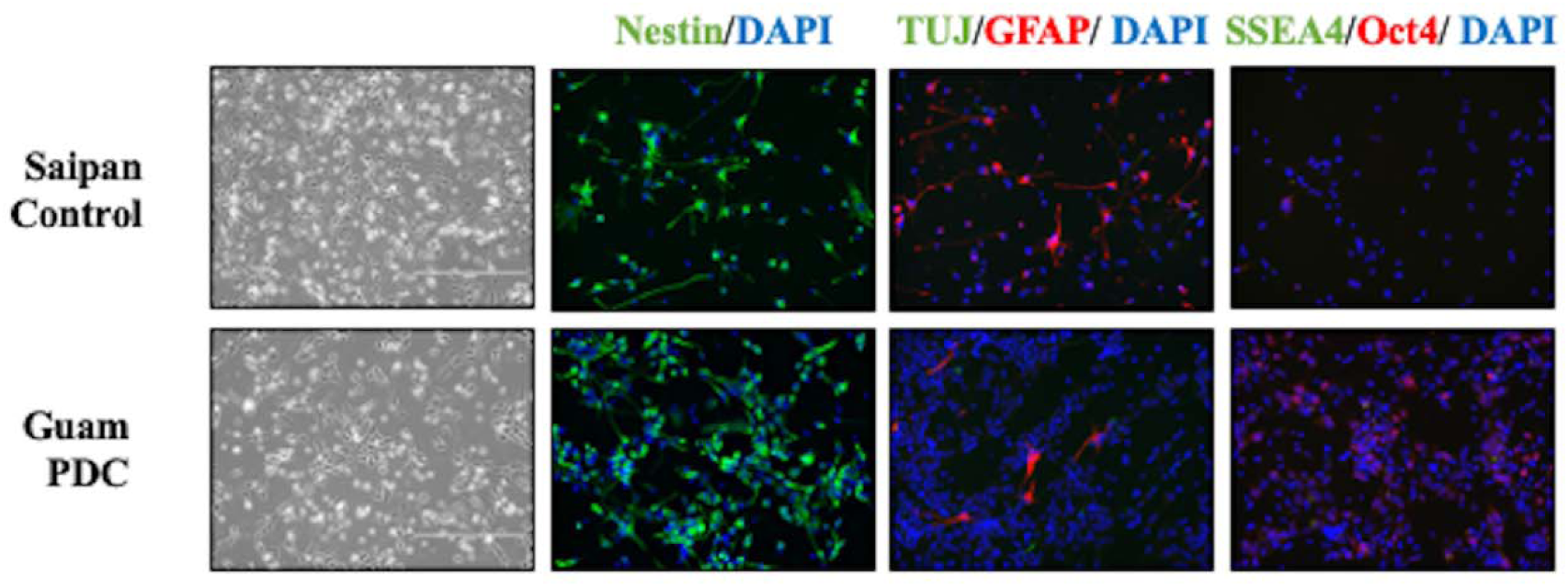
Characterization of Chamorro-derived astrocyte progenitor cells (APCs) APCs were differentiated from hNPCs using the STEMdiff^Tm^ protocol and mmunostained for astrocyte markers. Both Chamorro-derived lines were positive for the astrocyte marker GFAP and negative for pluripotency (SSEA4, OCT4) and neuronal differentiation (β−III-tubulin, TUJ) markers. Note that GFAP staining was more prominent in the Saipan control than the Guam PDC line. Cells were counterstained with DAPI nuclear stain.

The differentiation of astrocyte progenitors produced a cell population expressing S100β, GFAP and GLAST (**Figure 6**). S100B and GLAST are considered markers of mature astrocytes (Lattke and Guillemot 2022). The expression of vimentin and AQP4 is also evidence of the differentiation of both lines to an astrocytic lineage. The lower expression of these markers in the PDC line suggested that these cells were also less mature than those of the control cells.

**Figure 6.**
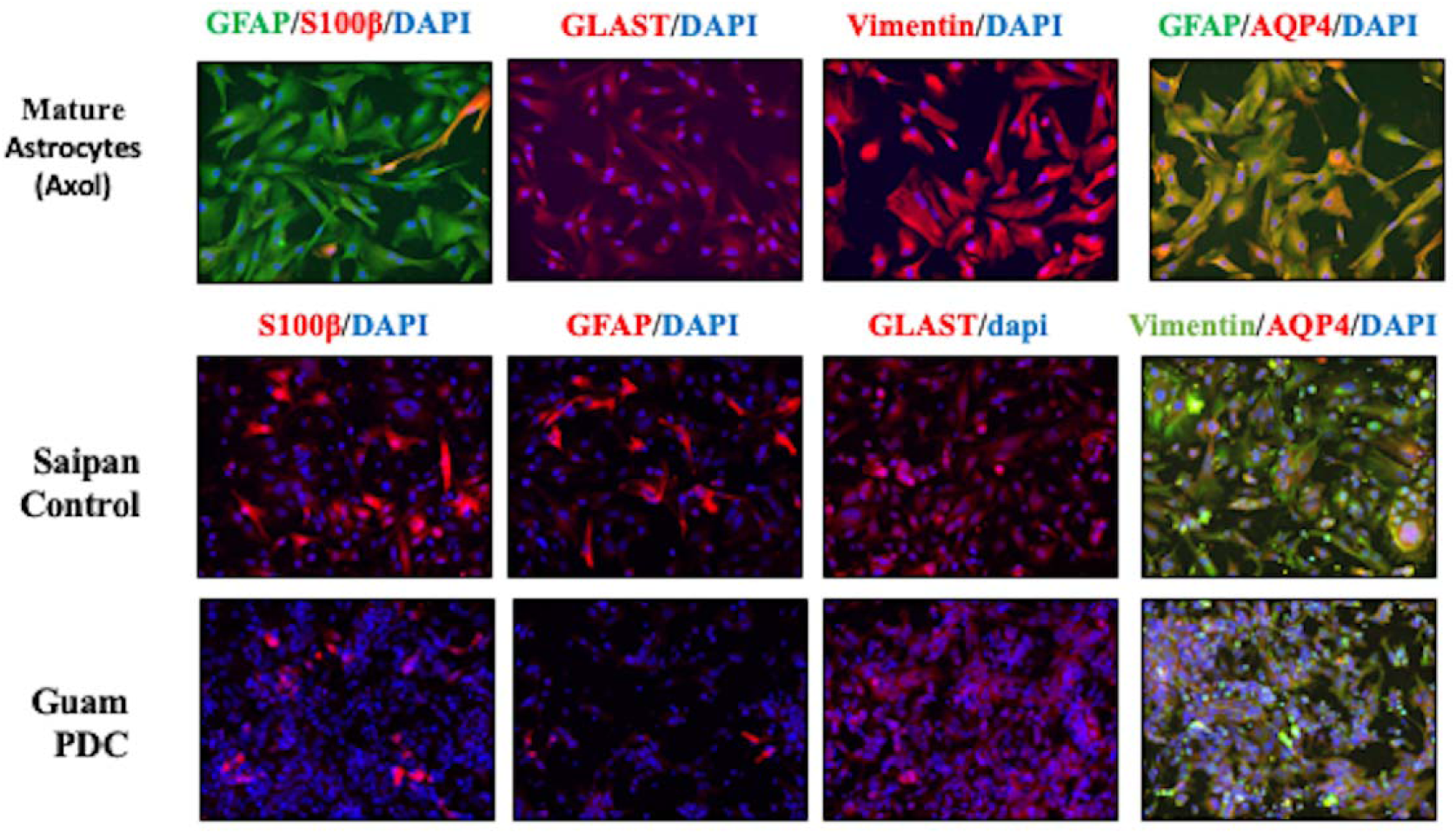
Characterization of Chamorro-derived mature astrocytes. Mature astrocytes from a manufacturer (Axol, *top images*) were compared with those derived from the hNPCs of the Chamorro patients. Mature astrocytes from Axol and both patients were immunostained for mature and other astrocyte markers (*bottom images*). Note that staining was more prominent in control than PDC astrocytes. Cells were counterstained with DAPI nucler stain. Abbreviations: S100b (S100 calcium-binding protein b), GLAST (glutamate aspartate transporter), and AQP4 (aquaporin 4).

Morphologically, most cells derived from the Saipan control and commercial mature astrocytes were flat and polygonal in shape in comparison to those from the PDC line. The reduced expression of astrocyte markers and the less complex morphology of PDC astrocytes suggested they were less mature than Saipan astrocytes. The development of astrocytes from additional patients will be required to confirm these findings.

## Discussion

Several contributing factors have been proposed for the etiology of Western Pacific ALS-PDC, including genetics (McGeer et al. 1997; Sieh et al. 2009; Spencer et al. 2016), infectious agents, notably a slow virus (prion) (Ahlskog et al. 1997; Condello et al. 2023; Gibbs and Gajdusek 1972), mineral imbalance (Spencer et al. 2005) and environmental toxins, notably those found in the *Cycas* plant, which was used traditionally as food and/or medicine in all three ALS/PDC disease foci (Spencer 1987; Spencer et al. 1987a; Spencer et al. 1987b; Spencer et al. 2016). Evidence in particular supports early-life exposure to the toxins in cycad seed that can lead to development of Western Pacific ALS/PDC later in life (Borenstein et al. 2007; Kisby and Spencer 2011); indeed some Guam and Kii-Japan cases show evidence of developmental cerebellar and retinal dysplasia attributable to *in utero* exposure during the second trimester to the active metabolite (methylazoxymethanol) of the principal cycad toxin (cycasin) (Spencer 2020). The availablility of human neural stem cells and neurons from age- and sex-matched non-neurological controls and from those with Western Pacific ALS/PDC should facilitate resolution of the etiology of this disorder like that for related neurodegenerative diseases (Beghini et al. 2024; Ceccarelli et al. 2024; Evangelisti et al. 2024; Israel et al. 2012). Emerging evidence indicates that patient-derived astrocytes also likely play a significant role in the pathogenesis of ALS/PDC in the Kii Peninsula of Japan (Kobayashi et al. 2023; Leventoux et al. 2024), which may also display evidence of developmental cerebellar and retinal perturbation consistent with cycad toxin exposure in utero (Spencer, 2020).

The ability to generate iPSCs from somatic cells has provided unprecedented opportunities for the study of neurodegenerative diseases where primary human tissues are not readily available (Evangelisti et al. 2024; Galgani et al. 2024). Furthermore, pathophysiology of specifically affected cell types can be gleaned from the differentiaton of iPSCs into different neural cell types (Lee et al. 2024; Yan et al. 2023). iPSCs were initially developed from dermal fibroblasts (Yu 2007, Takahashi 2007), but other somatic tissues like lymphoblastoid cells (LCLs) have gained importance (Fujimori et al. 2016; Gurwitz and Steeg 2024; Kim et al. 2023; Walker et al. 2021). LCLs have been reprogrammed into iPSCs for the differentiation to neuroprogenitor cells (hNPCs) and neurons as an *in vitro* model of neurodegenerative diseases (Fujimori et al. 2016; Hedges et al. 2021; Jung et al. 2023) that are functionally the same as fibroblast-derived iPSCs (Barret 2014, Corti 2015, Lee 2017, Shi 2012, Israel 2012, Verheyen 2015, Sareen 2012).

A major goal of these studies was to develop hiPSCs from LCLs of age- and sex-matched neurologically normal and PDC-affected Chamorros to explore the role of environmental factors in the etiology of Western Pacific ALS/PDC. iPSCs were developed from LCLs of Chamorro subjects, specifically a healthy control from Saipan and a patient with PDC from Guam. Although dermal fibroblasts are predominantly used for deriving iPSCs (Takahashi et al. 2007; Yu et al. 2007), skin tissue from Chamorros with and without PDC are not readily available. Furthermore, PDC has essentially disappeared on Guam and has been declining in Kii-Japan and Papua, Indonesia (Kuzuhara 2007; Okumiya et al. 2014; Spencer et al. 2005; Yoshida et al. 1998).

We have demonstrated that LCLs from Chamorro patients can be successfully reprogrammed into iPSCs and then differentiated into neural stem cells, neurons and astrocytes, as previously demonstrated for other neurodegenerative diseases (Barrett et al. 2014; Corti et al. 2015; Fujimori et al. 2016; Iovino et al. 2015; Israel et al. 2012; Kumar et al. 2016; Qu et al. 2023). A similar embroid body protocol has been used to develop immature and mature astrocytes from iPSCs of Kii-Japan ALS/PDC patients (Kobayashi et al. 2023; Leventoux et al. 2020). –The model system described here should be particulary useful for exploring the effects of environmental toxins in immature and mature neural stem cells derived from Guam PDC patients.

iPSCs from both PDC-affected and healthy Chamorros expressed all standard markers of pluripotency previously observed in iPSCs derived from fibroblasts or lymphoblastoid cell lines (Barrett et al. 2014; Fujimori et al. 2016), and they lost their pluripotency after their induction into neuroprogenitor cells (hNPCs). iPSCs from both patients also formed discrete colonies of tightly-packed cells (Maherali and Hochedlinger 2008). However, iPSCs from the control Saipan subject formed discrete colonies with well-defined edges whereas iPSCs from the Guam PDC subject grew more slowly with ragged edges. Similar morphological differences have been observed in iPSC lines derived from Kii-Japan ALS/PDC subjects (Kobayashi et al. 2023). Moreover, neither the LCL nor the iPSCs lines had chromosomal abnormalities, indicating the presence of a normal karyotype. Since the reprogramming of LCLs from both subjects was identical, the slower growth of iPSCs derived from the PDC patient might be due to genetic differences, as previously reported for some Guam PDC patients (Morris et al. 2004; Steele et al. 2015). Steele and colleagues (Steele et al. 2015) used targeting sequencing to identify 32 genes previously linked with Parkinson disease, dementia or ALS. Among the 32 PDC patients they examined, ten had mutations in disease-related genes (e.g., DJ-1, PINK1, SNCA). However, several studies have failed to demonstrate a consistent association between ALS-PDC and any neurodegenerative disease-associated genetic mutations (Figlewicz et al. 1994; Hermosura et al. 2008; Kaji et al. 2012; Kuzuhara and Kokubo 2005). The development of iPSCs from the other matched pairs of LCL lines (n=9) from both healthy control and PDC patient will be required to confirm our findings.

Our next goal was to determine if the hNPCs derived from both Chamorro iPSC lines differentiated into neural cells (i.e., neurons, astrocytes). hNPCs from both lines exhibited similar growth patterns and morphology (e.g., rosettes) to that of hNPCs derived from the reference iPSC line (*Supplement 2*). hNPCs from both iPSC lines also expressed neuroprogenitor markers (i.e., nestin), but they were negative for both pluripotency and neuronal markers. These studies demonstrate that hNPCs derived from both Chamorro patients provide an abundant source of neural stem cells for examining the effects of environmental factors on their growth and differentiation into various neural cells (Evangelisti et al. 2024).

Our final objective was to generate neurons and astrocytes from both hNPCs lines. The morphology of immature neurons (3 week-old) derived from both healthy control and PDC-affected hNPC lines exhibited similar expression of neural markers (nestin, Map2, DCX, and GFAP), but when they were allowed to mature for a longer period (i.e., 4-weeks) significant differences were noted in the expression of neuronal proteins (Map2, beta-tubulin III) as well as synaptic proteins (synaptophysin). PDC neurons that had been allowed to mature for a longer period showed a higher expression of nestin and lower expression of both microtubule proteins (TUJ and MAP2) and the synaptic protein synaptophysin. The expression of astrocyte markers also differed between the control and PDC-affected Chamorro subjects. Mature PDC astrocytes expressed lower levels of the calcium-binding protein S100b, a glutamate transporter (GLAST) and other markers of astrocytes (i.e, GFAP, Vimentin), which is consistent with differences noted in iPSC-derived astrocytes from Kii-Japan ALS/PDC patients (Kobayashi et al. 2023). These data suggest that the neurons of PDC patients were more immature than those from the healthy control, a finding that is consistent with that recently found for neurons derived from patients with AD (Mertens et al. 2021). Mertens and colleagues (Mertens et al. 2021) reported that neurons derived from AD patients develop a hypo-mature neuronal state characterized by markers of stress, cell cycle, and de-differentiation that was associated with epigenetic erosion (Herdy et al. 2022; Mertens et al. 2021). Jones and colleagues also found that astrocytes in both familial and sporadic AD patients exhibited a less complex morphological appearance, overall atrophic profiles and abnormal localization of functional astroglial markers (Jones et al. 2017). Such mechanisms might explain why PDC neurons and astrocytes derived from a Guam Chamorro PDC patient also exhibited a less complex morphology and a more immature phenotype than those derived from the healthy Saipan Chamorro subject, especially when they were allowed to mature for a longer period of time (Aldridge and West 2024). Comparable studies with the other age- and sex-matched pairs of iPSCs derived from LCLs control and Guam PDC patients will be required to confirm these findings.

## Conclusion

Neural stem cells produced from patient-derived iPSCs are proving to be a powerful model system for investigating the underlying pathogenesis of neurodegenerative diseases and neurodevelopmental disorders (Ardhanareeswaran et al., 2017; Efthymiou et al., 2014; Majolo et al., 2018; Mehta et al., 2018; Sareen et al., 2013). We demonstrated that iPSCs can also be developed from lymphoblastoid cell lines (LCLs) of healthy and PDC-impacted Chamorros for the purpose of investigating the underlying etiopathogenesis of Western Pacific ALS-PDC. We also identified differences in the differentiation of neurons and astrocytes from brain-diseased vs. healthy Chamorros that could be due to differences in environmental exposures and/or genetics. The heterogeneity at the iPSC stage is mainly driven by the genetic background of the donor rather than by any other non-genetic factor, such as culture conditions, passage or sex (Volpato and Webber 2020). Development and characterization of iPSCs, neurons and astrocytes from additional LCLs of age- and gender matched pairs of healthy Chamorro controls and PDC patients will clarify the source of these differences. Since Guam and Kii-Japan PDC are most likely triggered by early-in-life exposure to environmental agents(s) (Borenstein et al., 2007; Spencer et al., 2020), the development of neurons and astrocytes from the healthy Saipan Chamorro will provide important insight into the role of various environmental triggers of this disease, notably cycad toxins (e.g., cycasin/MAM, L-BMAA), their metabolites, and other agents (Spencer et al. 2020). Such studies should clarify the role of exposure to environmental triggers of Guam PDC.

## Supporting information

Supolementary Materials

## Acknowledgements

The authors would like to Dr Lousia Hooven and the CGRB at Oregon State University for granting access to the Keyence BZ-X710 fluorescent microscope for imaging. Supported by NIH 1R41ES026225-01 and an Intramural Grant from WUHS.

## Notes

### Competing Interest Statement

The authors have declared no competing interest.

### Summary of Updates

Acknowledgements about the source of the flourecent microscope used to obtain images and the funding support. This was inserted between the Conclusion and References sections.

## References

Ahlskog JE, Petersen RC, Waring SC, Esteban-Santillan C, Craig UK, Maraganore DM, Lennon VA, Kurland LT. 1997. Guamanian neurodegenerative disease: Are diabetes mellitus and altered humoral immunity clues to pathogenesis? Neurology. 48(5):1356–1362.

Aldridge AI, West AE. 2024. Epigenetics and the timing of neuronal differentiation. Curr Opin Neurobiol. 89:102915.

Amponsah AE, Guo R, Kong D, Feng B, He J, Zhang W, Liu X, Du X, Ma Z, Liu B et al. 2021. Patient-derived ipscs, a reliable in vitro model for the investigation of alzheimer’s disease. Rev Neurosci. 32(4):379–402.

Barrett R, Ornelas L, Yeager N, Mandefro B, Sahabian A, Lenaeus L, Targan SR, Svendsen CN, Sareen D. 2014. Reliable generation of induced pluripotent stem cells from human lymphoblastoid cell lines. Stem Cells Transl Med. 3(12):1429–1434.

Beghini DG, Kasai-Brunswick TH, Henriques-Pons A. 2024. Induced pluripotent stem cells in drug discovery and neurodegenerative disease modelling. Int J Mol Sci. 25(4).

Borenstein AR, Mortimer JA, Schofield E, Wu Y, Salmon DP, Gamst A, Olichney J, Thal LJ, Silbert L, Kaye J et al. 2007. Cycad exposure and risk of dementia, mci, and pdc in the chamorro population of guam. Neurology. 68(21):1764–1771.

Ceccarelli L, Verriello L, Pauletto G, Valente M, Spadea L, Salati C, Zeppieri M, Ius T. 2024. The role of human pluripotent stem cells in amyotrophic lateral sclerosis: From biological mechanism to practical implications. Front Biosci (Landmark Ed). 29(3):114.

Chapotte-Baldacci CA, Jauvin D, Chahine M. 2023. Generation of control ipsc lines cbrculi008-a and cbrculi009-a derived from lymphoblastoid cell lines. Stem Cell Res. 71:103168.

Chlebowski AC, Kisby GE. 2020. Protocol for high-throughput screening of neural cell or brain tissue protein using a dot-blot technique with near-infrared imaging. STAR Protoc. 1(2).

Choi SH, Kim YH, D’Avanzo C, Aronson J, Tanzi RE, Kim DY. 2015. Recapitulating amyloid beta and tau pathology in human neural cell culture models: Clinical implications. US Neurol. 11(2):102–105.

Choi SH, Kim YH, Hebisch M, Sliwinski C, Lee S, D’Avanzo C, Chen H, Hooli B, Asselin C, Muffat J et al. 2014. A three-dimensional human neural cell culture model of alzheimer’s disease. Nature. 515(7526):274–278.

Choi SH, Kim YH, Quinti L, Tanzi RE, Kim DY. 2016. 3d culture models of alzheimer’s disease: A road map to a “cure-in-a-dish”. Mol Neurodegener. 11(1):75.

Condello C, Ayers JI, Dalgard CL, Garcia Garcia MM, Rivera BM, Seeley WW, Perl DP, Prusiner SB. 2023. Guam als-pdc is a distinct double-prion disorder featuring both tau and abeta prions. Proc Natl Acad Sci U S A. 120(13):e2220984120.

Corti S, Faravelli I, Cardano M, Conti L. 2015. Human pluripotent stem cells as tools for neurodegenerative and neurodevelopmental disease modeling and drug discovery. Expert Opin Drug Discov. 10(6):615–629.

Evangelisti C, Ramadan S, Orlacchio A, Panza E. 2024. Experimental cell models for investigating neurodegenerative diseases. Int J Mol Sci. 25(17).

Figlewicz DA, Garruto RM, Krizus A, Yanagihara R, Rouleau GA. 1994. The cu/zn superoxide dismutase gene in als and parkinsonism-dementia of guam. Neuroreport. 5(5):557–560.

Forman MS, Schmidt ML, Kasturi S, Perl DP, Lee VM, Trojanowski JQ. 2002. Tau and alpha-synuclein pathology in amygdala of parkinsonism-dementia complex patients of guam. Am J Pathol. 160(5):1725–1731.

Fujimori K, Tezuka T, Ishiura H, Mitsui J, Doi K, Yoshimura J, Tada H, Matsumoto T, Isoda M, Hashimoto R et al. 2016. Modeling neurological diseases with induced pluripotent cells reprogrammed from immortalized lymphoblastoid cell lines. Mol Brain. 9(1):88.

Galgani A, Scotto M, Giorgi FS. 2024. The neuroanatomy of induced pluripotent stem cells: In vitro models of subcortical nuclei in neurodegenerative disorders. Curr Issues Mol Biol. 46(9):10180–10199.

Gibbs CJ, Jr., Gajdusek DC. 1972. Amyotrophic lateral sclerosis, parkinson’s disease, and the amyotrophic lateral sclerosis-parkinsonism-dementia complex on guam: A review and summary of attempts to demonstrate infection as the aetiology. J Clin Pathol Suppl (R Coll Pathol). 6:132–140.

Gimenez-Roldan S, Steele JC, Palmer VS, Spencer PS. 2021. Lytico-bodig in guam: Historical links between diet and illness during and after spanish colonization. J Hist Neurosci. 30(4):335–374.

Gurwitz D, Steeg R. 2024. Enriching ipsc research diversity: Harnessing human biobank collections for improved ethnic representation. Drug Dev Res. 85(5):e22227.

Hedges EC, Topp S, Shaw CE, Nishimura AL. 2021. Generation of six induced pluripotent stem cell lines from patients with amyotrophic lateral sclerosis with associated genetic mutations in either fus or anxa11. Stem Cell Res. 52:102246.

Herdy JR, Traxler L, Agarwal RK, Karbacher L, Schlachetzki JCM, Boehnke L, Zangwill D, Galasko D, Glass CK, Mertens J et al. 2022. Increased post-mitotic senescence in aged human neurons is a pathological feature of alzheimer’s disease. Cell Stem Cell. 29(12):1637–1652 e1636.

Hermosura MC, Cui AM, Go RC, Davenport B, Shetler CM, Heizer JW, Schmitz C, Mocz G, Garruto RM, Perraud AL. 2008. Altered functional properties of a trpm2 variant in guamanian als and pd. Proc Natl Acad Sci U S A. 105(46):18029–18034.

Iovino M, Agathou S, Gonzalez-Rueda A, Del Castillo Velasco-Herrera M, Borroni B, Alberici A, Lynch T, O’Dowd S, Geti I, Gaffney D et al. 2015. Early maturation and distinct tau pathology in induced pluripotent stem cell-derived neurons from patients with mapt mutations. Brain. 138(Pt 11):3345–3359.

Israel MA, Yuan SH, Bardy C, Reyna SM, Mu Y, Herrera C, Hefferan MP, Van Gorp S, Nazor KL, Boscolo FS, et al. 2012. Probing sporadic and familial alzheimer’s disease using induced pluripotent stem cells. Nature. 482(7384):216–220.

Jones VC, Atkinson-Dell R, Verkhratsky A, Mohamet L. 2017. Aberrant ipsc-derived human astrocytes in alzheimer’s disease. Cell Death Dis. 8(3):e2696.

Jung M, Hartmann C, Ehrhardt T, Peter LM, Abid CL, Harwardt B, Hirschfeld J, Claus C, Haferkamp U, Pless O et al. 2023. Generation of a set of induced pluripotent stem cell lines from two alzheimer disease patients carrying apoe4 (mlui007-j; mlui008-a) and healthy old donors carrying apoe3 (mlui009-a; mlui010-b) to study apoe in aging and disease. Stem Cell Res. 69:103072.

Kaji R, Izumi Y, Adachi Y, Kuzuhara S. 2012. Als-parkinsonism-dementia complex of kii and other related diseases in japan. Parkinsonism Relat Disord. 18 Suppl 1:S190–191.

Karmirian K, Holubiec M, Goto-Silva L, Fernandez Bessone I, Vitoria G, Mello B, Alloatti M, Vanderborght B, Falzone TL, Rehen S. 2023. Modeling alzheimer’s disease using human brain organoids. Methods Mol Biol. 2561:135–158.

Kim M, Park J, Kim S, Han DW, Shin B, Scholer HR, Kim J, Kim KP. 2023. Generation of induced pluripotent stem cells from lymphoblastoid cell lines by electroporation of episomal vectors. Int J Stem Cells. 16(1):36–43.

Kim Y, Yun B, Ye BS, Kim BY. 2024. Generation of alzheimer’s disease model derived from induced pluripotent stem cells with app gene mutation. Biomedicines. 12(6).

Kisby GE, Spencer PS. 2011. Is neurodegenerative disease a long-latency response to early-life genotoxin exposure? Int J Environ Res Public Health. 8(10):3889–3921.

Kobayashi H, Ueda K, Morimoto S, Ishikawa M, Leventoux N, Sasaki R, Hirokawa Y, Kokubo Y, Okano H. 2023. Protein profiling of extracellular vesicles from ipsc-derived astrocytes of patients with als/pdc in kii peninsula. Neurol Sci. 44(12):4511–4516.

Kumar S, Curran JE, Espinosa EC, Glahn DC, Blangero J. 2020. Highly efficient induced pluripotent stem cell reprogramming of cryopreserved lymphoblastoid cell lines. J Biol Methods. 7(1):e124.

Kumar S, Curran JE, Glahn DC, Blangero J. 2016. Utility of lymphoblastoid cell lines for induced pluripotent stem cell generation. Stem Cells Int. 2016:2349261.

Kuzuhara S. 2007. [muro disease or als-parkinsonism-dementia complex of the kii peninsula of japan]. Rinsho Shinkeigaku. 47(11):695–702.

Kuzuhara S, Kokubo Y. 2005. Atypical parkinsonism of japan: Amyotrophic lateral sclerosis-parkinsonism-dementia complex of the kii peninsula of japan (muro disease): An update. Mov Disord. 20 Suppl 12:S108–113.

Lattke M, Guillemot F. 2022. Understanding astrocyte differentiation: Clinical relevance, technical challenges, and new opportunities in the omics era. WIREs Mech Dis. 14(5):e1557.

Lee DH, Lee EC, Lee JY, Lee MR, Shim JW, Oh JS. 2024. Neuronal cell differentiation of ipscs for the clinical treatment of neurological diseases. Biomedicines. 12(6).

Lee H, Pearse RV, 2nd, Lish AM, Pan C, Augur ZM, Terzioglu G, Gaur P, Liao M, Fujita M, Tio ES et al. 2025. Contributions of genetic variation in astrocytes to cell and molecular mechanisms of risk and resilience to late-onset alzheimer’s disease. Glia. 73(6):1166–1187.

Leventoux N, Morimoto S, Imaizumi K, Sato Y, Takahashi S, Mashima K, Ishikawa M, Sonn I, Kondo T, Watanabe H et al. 2020. Human astrocytes model derived from induced pluripotent stem cells. Cells. 9(12).

Leventoux N, Morimoto S, Ishikawa M, Nakamura S, Ozawa F, Kobayashi R, Watanabe H, Supakul S, Okamoto S, Zhou Z et al. 2024. Aberrant chchd2-associated mitochondriopathy in kii als/pdc astrocytes. Acta Neuropathol. 147(1):84.

Li L, Chao J, Shi Y. 2018. Modeling neurological diseases using ipsc-derived neural cells : Ipsc modeling of neurological diseases. Cell Tissue Res. 371(1):143–151.

MacArthur CC, Pradhan S, Wetton N, Zarrabi A, Dargitz C, Sridharan M, Jackson S, Pickle L, Lakshmipathy U. 2019. Generation and comprehensive characterization of induced pluripotent stem cells for translational research. Regen Med. 14(6):505–524.

Maherali N, Hochedlinger K. 2008. Guidelines and techniques for the generation of induced pluripotent stem cells. Cell Stem Cell. 3(6):595–605.

Malik N, Wang X, Shah S, Efthymiou AG, Yan B, Heman-Ackah S, Zhan M, Rao M. 2014. Comparison of the gene expression profiles of human fetal cortical astrocytes with pluripotent stem cell derived neural stem cells identifies human astrocyte markers and signaling pathways and transcription factors active in human astrocytes. PLoS One. 9(5):e96139–e96139.

Marei HE, Khan MUA, Hasan A. 2023. Potential use of ipscs for disease modeling, drug screening, and cell-based therapy for alzheimer’s disease. Cell Mol Biol Lett. 28(1):98.

McGeer PL, Schwab C, McGeer EG, Haddock RL, Steele JC. 1997. Familial nature and continuing morbidity of the amyotrophic lateral sclerosis-parkinsonism dementia complex of guam. Neurology. 49(2):400–409.

McGeer PL, Steele JC. 2011. The als/pdc syndrome of guam: Potential biomarkers for an enigmatic disorder. Prog Neurobiol. 95(4):663–669.

Mertens J, Herdy JR, Traxler L, Schafer ST, Schlachetzki JCM, Bohnke L, Reid DA, Lee H, Zangwill D, Fernandes DP et al. 2021. Age-dependent instability of mature neuronal fate in induced neurons from alzheimer’s patients. Cell Stem Cell. 28(9):1533–1548 e1536.

Miklossy J, Steele JC, Yu S, McCall S, Sandberg G, McGeer EG, McGeer PL. 2008. Enduring involvement of tau, beta-amyloid, alpha-synuclein, ubiquitin and tdp-43 pathology in the amyotrophic lateral sclerosis/parkinsonism-dementia complex of guam (als/pdc). Acta Neuropathol. 116(6):625–637.

Mimuro M, Yoshida M, Kuzuhara S, Kokubo Y. 2018. Amyotrophic lateral sclerosis and parkinsonism-dementia complex of the hohara focus of the kii peninsula: A multiple proteinopathy? Neuropathology. 38(1):98–107.

Morris HR, Steele JC, Crook R, Wavrant-De Vrieze F, Onstead-Cardinale L, Gwinn-Hardy K, Wood NW, Farrer M, Lees AJ, McGeer PL et al. 2004. Genome-wide analysis of the parkinsonism-dementia complex of guam. Arch Neurol. 61(12):1889–1897.

Okano H, Morimoto S. 2022. Ipsc-based disease modeling and drug discovery in cardinal neurodegenerative disorders. Cell Stem Cell. 29(2):189–208.

Okumiya K, Wada T, Fujisawa M, Ishine M, Garcia Del Saz E, Hirata Y, Kuzuhara S, Kokubo Y, Seguchi H, Sakamoto R et al. 2014. Amyotrophic lateral sclerosis and parkinsonism in papua, indonesia: 2001-2012 survey results. BMJ Open. 4(4):e004353.

Pandya H, Shen MJ, Ichikawa DM, Sedlock AB, Choi Y, Johnson KR, Kim G, Brown MA, Elkahloun AG, Maric D et al. 2017. Differentiation of human and murine induced pluripotent stem cells to microglia-like cells. Nature Neuroscience. 20(5):753–759.

Pazzin DB, Previato TTR, Budelon Goncalves JI, Zanirati G, Xavier FAC, da Costa JC, Marinowic DR. 2024. Induced pluripotent stem cells and organoids in advancing neuropathology research and therapies. Cells. 13(9).

Qu W, Canoll P, Hargus G. 2023. Molecular insights into cell type-specific roles in alzheimer’s disease: Human induced pluripotent stem cell-based disease modelling. Neuroscience. 518:10–26.

Rajesh D, Dickerson SJ, Yu J, Brown ME, Thomson JA, Seay NJ. 2011. Human lymphoblastoid b-cell lines reprogrammed to ebv-free induced pluripotent stem cells. Blood. 118(7):1797–1800.

Ramme AP, Faust D, Koenig L, Nguyen N, Marx U. 2021. Supporting dataset of two integration-free induced pluripotent stem cell lines from related human donors. Data Brief. 37:107140.

Riemens RJM, Kenis G, van den Beucken T. 2020. Human-induced pluripotent stem cells as a model for studying sporadic alzheimer’s disease. Neurobiol Learn Mem. 175:107318.

Rivetti di Val Cervo P, Besusso D, Conforti P, Cattaneo E. 2021. Hipscs for predictive modelling of neurodegenerative diseases: Dreaming the possible. Nat Rev Neurol. 17(6):381–392.

Samson JS, Ramesh A, Parvathi VD. 2024. Development of midbrain dopaminergic neurons and the advantage of using hipscs as a model system to study parkinson’s disease. Neuroscience. 546:1–19.

Sebeo J, Hof PR, Perl DP. 2004. Occurrence of alpha-synuclein pathology in the cerebellum of guamanian patients with parkinsonism-dementia complex. Acta Neuropathol. 107(6):497–503.

Sie L, Loong S, Tan EK. 2009. Utility of lymphoblastoid cell lines. J Neurosci Res. 87(9):1953–1959.

Sieh W, Choi Y, Chapman NH, Craig UK, Steinbart EJ, Rothstein JH, Oyanagi K, Garruto RM, Bird TD, Galasko DR et al. 2009. Identification of novel susceptibility loci for guam neurodegenerative disease: Challenges of genome scans in genetic isolates. Hum Mol Genet. 18(19):3725–3738.

Smith R, Hovren H, Bowser R, Bakkar N, Garruto R, Ludolph A, Ravits J, Gaertner L, Murphy D, Lebovitz R. 2024. Misfolded alpha-synuclein in amyotrophic lateral sclerosis: Implications for diagnosis and treatment. Eur J Neurol. 31(4):e16206.

Spencer PS. 1987. Guam als/parkinsonism-dementia: A long-latency neurotoxic disorder caused by “slow toxin(s)” in food? Can J Neurol Sci. 14(3 Suppl):347–357.

Spencer PS. 2020. Etiology of retinal and cerebellar pathology in western pacific amyotrophic lateral sclerosis and parkinsonism-dementia complex. Eye Brain. 12:97–104.

Spencer PS. 2022. Parkinsonism and motor neuron disorders: Lessons from western pacific als/pdc. J Neurol Sci. 433:120021.

Spencer PS, Ohta M, Palmer VS. 1987a. Cycad use and motor neurone disease in kii peninsula of japan. Lancet. 2(8573):1462–1463.

Spencer PS, Palmer VS, Herman A, Asmedi A. 1987b. Cycad use and motor neurone disease in irian jaya. Lancet. 2(8570):1273–1274.

Spencer PS, Palmer VS, Kisby GE. 2016. Seeking environmental causes of neurodegenerative disease and envisioning primary prevention. Neurotoxicology. 56:269–283.

Spencer PS, Palmer VS, Kisby GE. 2020. Western pacific als-pdc: Evidence implicating cycad genotoxins. J Neurol Sci. 419:117185.

Spencer PS, Palmer VS, Ludolph AC. 2005. On the decline and etiology of high-incidence motor system disease in west papua (southwest new guinea). Mov Disord. 20 Suppl 12:S119–126.

Steele JC, Guella I, Szu-Tu C, Lin MK, Thompson C, Evans DM, Sherman HE, Vilarino-Guell C, Gwinn K, Morris H et al. 2015. Defining neurodegeneration on guam by targeted genomic sequencing. Ann Neurol. 77(3):458–468.

Takahashi K, Tanabe K, Ohnuki M, Narita M, Ichisaka T, Tomoda K, Yamanaka S. 2007. Induction of pluripotent stem cells from adult human fibroblasts by defined factors. Cell. 131(5):861–872.

Trevisan M, Desole G, Costanzi G, Lavezzo E, Palu G, Barzon L. 2017. Reprogramming methods do not affect gene expression profile of human induced pluripotent stem cells. Int J Mol Sci. 18(1).

Valadez-Barba V, Juarez-Navarro K, Padilla-Camberos E, Diaz NF, Guerra-Mora JR, Diaz-Martinez NE. 2021. Parkinson’s disease: An update on preclinical studies of induced pluripotent stem cells. Neurologia (Engl Ed).

Verheijen BM, Morimoto S, Sasaki R, Oyanagi K, Kokubo Y, Kuzuhara S, van Leeuwen FW. 2020. Expression of mutant ubiquitin and proteostasis impairment in kii amyotrophic lateral sclerosis/parkinsonism-dementia complex brains. J Neuropathol Exp Neurol. 79(8):902–907.

Volpato V, Webber C. 2020. Addressing variability in ipsc-derived models of human disease: Guidelines to promote reproducibility. Dis Model Mech. 13(1).

Walker SJ, Wagoner AL, Leavitt D, Mack DL. 2021. A simplified approach for derivation of induced pluripotent stem cells from epstein-barr virus immortalized b-lymphoblastoid cell lines. Heliyon. 7(4):e06617.

Wehr NB, Levine RL. 2012. Quantitation of protein carbonylation by dot blot. Anal Biochem. 423(2):241–245.

Winton MJ, Joyce S, Zhukareva V, Practico D, Perl DP, Galasko D, Craig U, Trojanowski JQ, Lee VM. 2006. Characterization of tau pathologies in gray and white matter of guam parkinsonism-dementia complex. Acta Neuropathol. 111(5):401–412.

Yan YW, Qian ES, Woodard LE, Bejoy J. 2023. Neural lineage differentiation of human pluripotent stem cells: Advances in disease modeling. World J Stem Cells. 15(6):530–547.

Yoshida S, Uebayashi Y, Kihira T, Kohmoto J, Wakayama I, Taguchi S, Yase Y. 1998. Epidemiology of motor neuron disease in the kii peninsula of japan, 1989-1993: Active or disappearing focus? J Neurol Sci. 155(2):146–155.

Yu J, Vodyanik MA, Smuga-Otto K, Antosiewicz-Bourget J, Frane JL, Tian S, Nie J, Jonsdottir GA, Ruotti V, Stewart R et al. 2007. Induced pluripotent stem cell lines derived from human somatic cells. Science. 318(5858):1917–1920.

Zhang ZX, Anderson DW, Mantel N. 1990. Geographic patterns of parkinsonism-dementia complex on guam. 1956 through 1985. Arch Neurol. 47(10):1069–1074.

