## Supplementary material for "Development of Patient-Derived Neuroprogenitor Cells (hNPCs), Neurons and Astrocytes to Explore the Etiology of Guam Parkinsonism-Dementia Complex (PDC)": Supolementary Materials

### Supplemental Materials

#### Supplement 1: Generation of iPSCs from LCLs.

**Table S1. Plasmids used for the transfection of LCLs.**

| Plasmid name | Addgene catalog number | Gene/insert name |
|---|---|---|
| pCXLE-hSK | 27078 | SOX2, KLF4 |
| pCXLE-hUL | 27080 | L-MYC |
| pCXLE-hOCT3/4-shp53-F | 27077 | OCT3/4 |
| pEP4 E02S ET2K | 20927 | OCT4, SOX2, SV40LT, Klf4 |

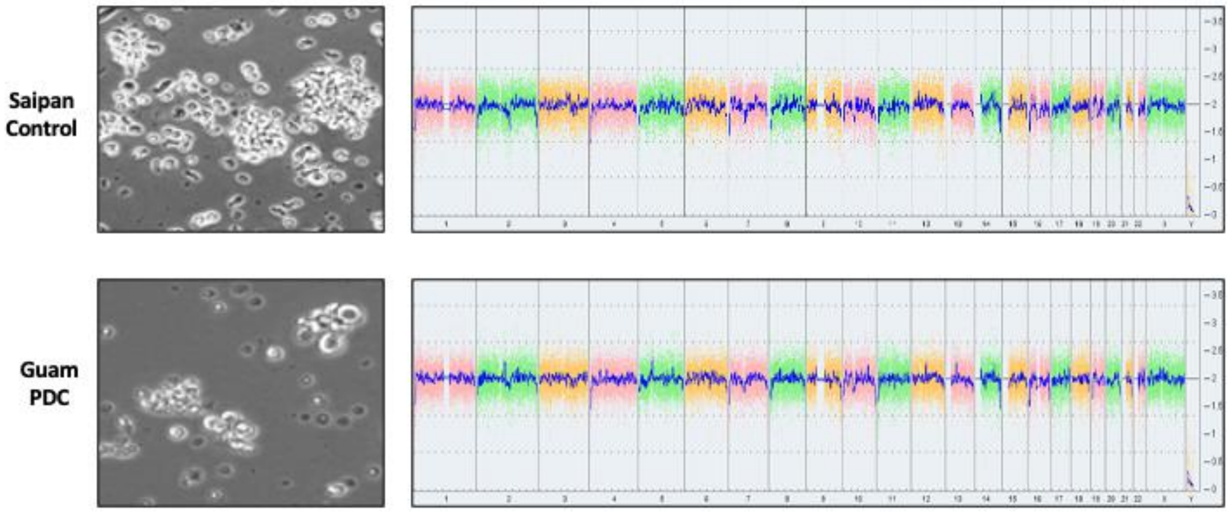


**Figure S1. Karyostat analysis of LCLs from healthy and PDC Chamorro patients**. The whole genome view displays all somatic and sex chromosomes in one frame with high level copy number. The smooth signal plot (right y-axis) is the smoothing of the log2 ratios which depict the signal intensities of probes on the microarray. A value of 2 represents a normal copy number state (CN = 2). A value of 3 represents chromosomal gain (CN = 3). A value of 1 represents a chromosomal loss (CN = 1). The pink, green and yellow colors indicate the raw signal for each individual chromosome probe, while the blue signal represents the normalized probe signal.

**Supplement 2**: **Optimization Method for the generation of hNPCs**

The reference iPSC line NCRM5 (ND50031) was obtained from RUCDR Infinite Biologics (Piscataway, NJ) and cultured on Matrigel® with mTeSR®-1 media. This line differentiates into hNPCs, neurons, and astrocytes (Malik et al., 2014, Pandya et al., 2017) and hNPCs derived from this line are commercially available from Axol Bioscience (Cambridge, UK).

For the generation of hNPCs, three commercially available methods were compared: The STEMdiff™ monolayer protocol with SMADi (STEMCELL™ Technologies), the STEMdiff™ Embryoid Body protocol with SMADi (STEMCELL™ Technologies), and the Cortical Neural Induction Kit ("Axol Monolayer", Axol Bioscience). NCRM5 iPSCs were cultured on Matrigel® with mTeSR®-1 media prior to starting either STEMdiff™ protocol, or were transitioned to vitronectin-coated plasticware and cultured in Essential 8™ Medium (ThermoFisher) for 2-3 passages before starting each protocol. All three protocols followed the manufacturer's instructions, except the Axol protocol added 20 µM PluriSln-1 (STEMCELL™ Technologies) on day 13 to the culture media and it was removed on day 15. This reportedly removed any iPSCs that remained in the culture (J. Crowe, Aston University, *personal communication*).

Following completion of the iPSC protocols, cells were cultivated in the appropriate NPC media (STEMdiff™ NPC medium or Axol Neural Expansion-XF Medium) for a minimum of two passages and their morphology monitored daily. Cells were fixed at each step for characterization by immunocytochemistry. Additionally, hNPCs at each step of the protocol were evaluated for the efficiency of passaging, cryopreservation and recovery. hNPCs from all three protocols displayed the expected cell morphology and nestin expression (*Figure S2*). hNPCs with the best morphology, nestin expression and efficiency of cultivation were those generated using the STEMdiff™ Embryoid Body protocol. This method was used for the differentiation of hNPCs into neural lineages (i.e., neurons, glia).

The STEMdiff™ Embryoid Body method consists of seeding $3x10^6$ iPSCs at passage 10+ on each well of an AggreWell™ 800 24-well plate (STEMCELL™ Technologies) containing Neural Induction Medium plus SMADi (STEMCELL™ Technologies) that was supplemented with 10 µM Y-27632 (STEMCELL™ Technologies) only on the first day of plating. Embryoid bodies (EBs) that developed were fed daily for five days before they were collected and transferred to a Matrigel®-coated plate. On the $8^{th}$ day, attached EBs were evaluated visually for the efficiency of neural induction (i.e., neural rosette formation) On the $12^{th}$ day, hNPCs were passaged (P0) onto Matrigel®-coated dishes, and then they passed again and were switched to STEMdiff™ Neural Progenitor Medium (STEMCELL™ Technologies) at day 17-19 (passage 1).

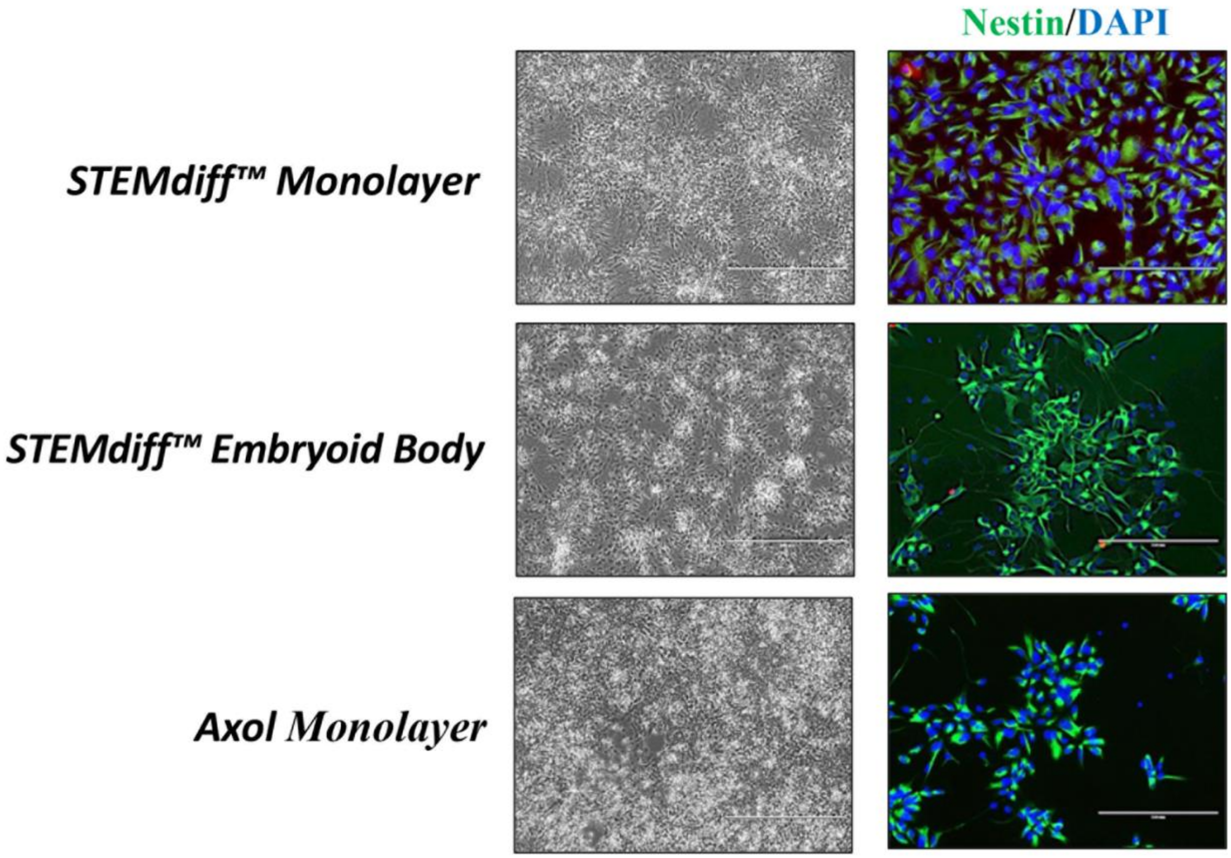


**Figure S2. Methods used to generate NPCs from a reference iPSC line.** The reference line NCRM5 (passage 20+) was used to generate hNPCs using three commercial protocols and the morphology and nestin staining compared by microscopy. Immunostaining for nestin (green, with dapi nuclear counterstain). Both STEMdiff™ kits utilized the SMADi as recommended by the manufacturer. Note that the STEMdiff™ Embryoid Body protocol gave the best morphology and nestin staining for the induction of the iPSC reference line.

**Supplement 3**: **Method used for the generation of neurons**

NCRM5-derived hNPCs using the STEMdiff™ Embryoid Body protocol were treated with three commercially available medias for their differentiation into neurons: Axol Neural Differentiation-XF Media (Axol Bioscience), Axol Neural Maintenance-XF Media (Axol Bioscience), and BrainPhys™ Neuronal Medium (STEMCELL™ Technologies). The two medias from Axol Bioscience were used as purchased. Supplements were added to the BrainPhys™ media to promote neuronal differentiation: 1x NeuroCult™ SM1 Supplement (STEMCELL™ Technologies), 1x N2 Supplement A (STEMCELL™ Technologies), 20 ng/mL BDNF (Invitrogen), 20 ng/mL GDNF (Peprotech), 1 mM dibutyryl c-AMP (STEMCELL™ Technologies), 200 nM ascorbic acid (STEMCELL™ Technologies). The hNPCs were differentiated for 21 days according to the manufacturer's protocols and the cells evaluated for morphology, cell migration, and the expression of neuronal markers (Figure S3) (antibodies listed in listed in Supplement 4). The BrainPhys™ protocol produced immature neurons with minimal cell migration and the morphology and neuronal markers characteristic of immature neurons.   This method was used to generate neurons from Chamorro-derived hNPCs.

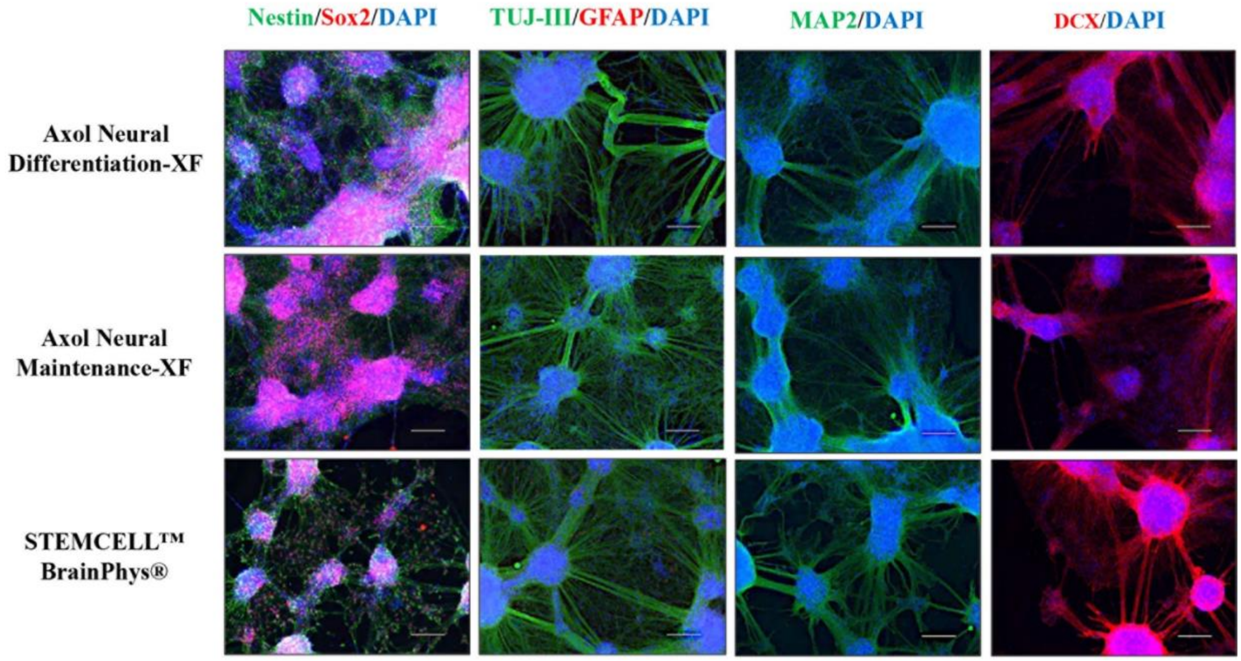


**Figure S3. Comparison of methods used to differentiate NCRM5-derived hNPCs into neurons.** Representative images of neurons immunoprobed with various markers after 21 days of differentiation. Note the staining of hNPCs (Nestin/Sox2) was very low whereas most cells were heavily stained for immature (TUJ-III, DCX) or mature (MAP2) neuronal markers with little/no staining for glial differentiation (GFAP). TUJ-III is β-tubulin III; MAP2 is microtubule associated protein 2; DCX is doublecortin; GFAP is glial fibrillary acidic protein.

**Supplement 4**: **Immunocytochemical characterization of iPSCs, hNPCs, neurons and astrocytes**

*iPSCs:* The Pluripotent Stem Cell 4-Marker Immunocytochemistry Kit (Invitrogen) was used for immunoprobing iPSCs. Following fixation with 4% paraformaldehyde for 20-30 minutes, cells were permeabilized for 15 min, blocked for 30 min and then incubated for 3 hr with antibodies against SSEA-4,OCT-4, SOX2, and TRA-1-60 and then for 1 hr with secondary antibodies at RT according to the manufacturer's protocol.  The immunoprobed cells were then counterstained with DAPI (NuncBlue®, Invitrogen) prior to imaging.

*hNPCs, neurons and astrocytes:* hNPCs, neurons and astrocytes that had been fixed with 4% paraformaldehyde for 20-30 minutes were blocked for 2 hr with buffer (5% BSA, 1.5% normal goat serum, 0.2% Triton-X in PBS), before incubation for 1 hr with primary antibodies and then for 2 hr with secondary antibodies at RT. Primary antibodies used for the characterization of NPCs, neurons astrocyte precursors or mature astrocytes are listed in Table S2. Secondary antibodies were AlexaFluor® 488 goat-anti-mouse and AlexaFluor® 568 goat-anti-rabbit (Invitrogen, 1:250). The immunoprobed cells were then counterstained with DAPI (NuncBlue®, Invitrogen) prior to imaging.

**Table S2. Antibodies used for Immunoprobing hNPCs, neurons and astrocytes**

| Antibody/Target | Source | Dilution factor | Species | Catalog Number |
|---|---|---|---|---|
| Nestin (10C2) | MilliporeSigma | 1:500 | Mouse | MA1-110 |
| Sox2 | MilliporeSigma | 1:500 | Rabbit | AB5603 |
| Beta-tubulin-III (TU-20) | MilliporeSigma | 1:1000 | Mouse | MAB1637 |
| GFAP | MilliporeSigma | 1:500 | Rabbit | AB5804 |
| MAP2 (AP18) | Invitrogen | 1:1000 | Mouse | MA5-12826 |
| DCX | Abcam | 1:1000 | Rabbit | ab18723 |
| Synaptophysin (D8F6H) | Cell Signaling | 1:1000 | Rabbit | 36406 |
| PSD-95 (D27E11) | Cell Signaling | 1:1000 | Rabbit | 3450 |
| Vimentin | Proteintech | 1:50 | Mouse | 60330-1-Ig |
| GLAST | Proteintech | 1:20 | Rabbit | 20785-1-AP |
| Aquaporin A (AQP4) | Proteintech | 1:100 | Rabbit | 16473-1-AP |
| S100β | Proteintech | 1:50 | Rabbit | 15146-1-AP |

**Supplement 5**: **Dot blot assay**

Details on this method have been previously published (Chlebowski and Kisby, 2020). Briefly, protein samples from both Saipan control and Guam PDC subjects (n=2/cell type) were thawed and diluted with TBS before application to pre-soaked nitrocellulose membranes (0.45 µm, BioRad) for dot blotting. The diluted samples (1 µg protein) were applied in duplicate to the membrane, air-dried for 1 hr and stored at 4°C prior to imaging. The membranes were first probed with Revert® 700 Total Protein Stain Kit (LiCor, Lincoln Nebraska) to determine equal loading of samples. The wetted membranes were incubated at RT on a rocker for 5 min with the Revert® stain, then washed 1x in Revert wash and distilled water prior to imaging on a near infrared imager (i.e., Odyssey CLx, LiCor). The membranes were then destained for 5-7 min at RT and then rinsed with distilled water. The destained membrane was then re-imaged to measure residual signal. The destained membrane was blocked for 1 hr in LiCor Intercept® (TBS) buffer, the membrane incubated with primary antibodies (Table S3) overnight at 4°C, and the next day the membrane washed in TBST (TBS +0.1% Tween-20) before application of the LiCor secondary antibody 1:10,000 in blocking buffer for 1 hr in the dark before they were imaged.

**Table S3: Antibodies used for dot blotting hNPCs and neurons**

| Antibody/Target | Source | Dilution factor | Species | Catalog Number |
|---|---|---|---|---|
| Nestin (10C2) | MilliporeSigma | 1:1000 | Mouse | MA1-110 |
| Beta-tubulin-III (TU-20) | MilliporeSigma | 1:1000 | Mouse | MAB1637 |
| GFAP | Proteintech | 1:5000 | Mouse | 60190-1-Ig |
| MAP2 (AP18) | Invitrogen | 1:1000 | Mouse | MA5-12826 |
| DCX | Abcam | 1:1000 | Rabbit | ab18723 |
| Tau5 | Invitrogen | 1:1000 | Mouse | MA5-12808 |
| Synaptophysin (D8F6H) | Cell Signaling | 1:1000 | Rabbit | 36406 |
| PSD-95 (D27E11) | Cell Signaling | 1:1000 | Rabbit | 3450 |

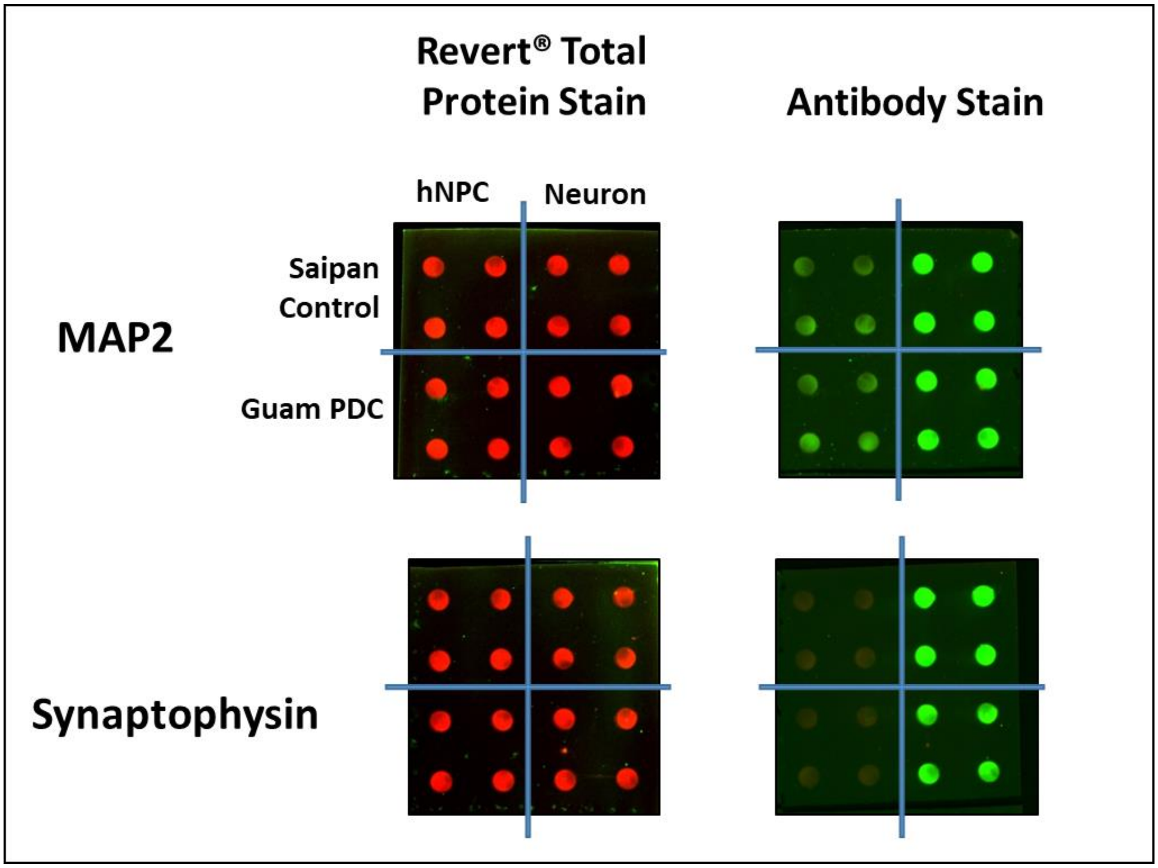


**Figure S4. Representative dot blot images of hNPCs and neurons**. Images were taken of dot blot membranes following staining with Revert® 700 Total Protein Stain or antibody incubations for either MAP2 (microtubule associated protein 2) or synaptophysin of the cell lysate source indicated (Saipan control or Guam PDC line, hNPC or neuron). Note the uniformity in the Revert® signals.

CHLEBOWSKI, A. C. & KISBY, G. E. 2020. Protocol for High-Throughput Screening of Neural Cell or Brain Tissue Protein Using a Dot-Blot Technique with Near-Infrared Imaging. *STAR Protocols,* 1**,** 100054.

MALIK, N., WANG, X., SHAH, S., EFTHYMIOU, A. G., YAN, B., HEMAN-ACKAH, S., ZHAN, M. & RAO, M. 2014. Comparison of the gene expression profiles of human fetal cortical astrocytes with pluripotent stem cell derived neural stem cells identifies human astrocyte markers and signaling pathways and transcription factors active in human astrocytes. *PloS one,* 9**,** e96139-e96139.

PANDYA, H., SHEN, M. J., ICHIKAWA, D. M., SEDLOCK, A. B., CHOI, Y., JOHNSON, K. R., KIM, G., BROWN, M. A., ELKAHLOUN, A. G., MARIC, D., SWEENEY, C. L., GOSSA, S., MALECH, H. L., MCGAVERN, D. B. & PARK, J. K. 2017. Differentiation of human and murine induced pluripotent stem cells to microglia-like cells. *Nature Neuroscience,* 20**,** 753-759.
